# MEF2D impairs mitochondrial respiration, glucose-stimulated insulin secretion, and survival in INS-1 β-cells

**DOI:** 10.64898/2026.03.07.709963

**Authors:** Jacqueline E. Crabtree, Rohit B. Sharma, Jeffery S. Tessem

## Abstract

Myocyte Enhancer Factor 2D (Mef2D) is a member of the Mef2 family. As a transcription factor, Mef2D regulates the expression of genes that impinge on cellular viability, tissue development, and fuel metabolism in a tissue dependent manner. Mef2D is expressed in the beta-cell, and overexpression and knockdown have been shown to modulate glucose stimulated insulin secretion. We sought to understand the role of Mef2D on beta-cell function and survival. To determine the function of Mef2D in the beta-cell, we built overexpression and knockdown INS-1 832/13 cell lines. We determined the effect of Mef2D overexpression or knockdown on mitochondrial respiration, insulin secretion, cell survival, and gene expression. Our data demonstrates that Mef2D knockdown enhances mitochondrial respiration, insulin secretion, and cell survival. Conversely, Mef2D overexpression inhibits mitochondrial respiration, insulin secretion, and cell survival. We demonstrate that some of this effect is due to modulated expression of the mitochondrial gene mtND6. These findings demonstrate that Mef2D overexpression is detrimental to beta-cell function and that Mef2D knockdown is beneficial. These data suggest that Mef2D may be a viable target to enhance functional beta-cell mass as a treatment for Type 1 and Type 2 Diabetes.

## INTRODUCTION

The pancreatic beta-cell controls glucose homeostasis by producing and secreting insulin in response to hyperglycemia. Insulin causes the muscle, liver, and adipose tissues to absorb and store glucose, thus causing the body to return to normoglycemia. Therefore, properly functioning beta-cell mass is essential for normal physiology. The incidence of Type 2 Diabetes (T2D) is increasing quite rapidly. As of 2018, 10.5% of the US population was diagnosed with diabetes.

The cost of treating those with diabetes in 2017 was $327 billion [1]. While T2D is associated with muscle, liver, and adipose insulin insensitivity, data from GWAS studies demonstrate that many genes associated with T2D are beta-cell-specific [2–6]. As such, patients with T2D demonstrate decreased functional beta-cell mass, defined by impaired insulin secretion, decreased cellular proliferation, and decreased survival. This change in function and survival is due to glucolipotoxic conditions that result in endoplasmic reticulum stress. Defining and understanding the molecular pathways that impinge on these phenotypes is necessary to design interventions to maintain or increase functional beta-cell mass to develop therapies to relieve the burden of T2D.

Myocyte enhancer factor 2D (Mef2D) is a transcription factor belonging to the MEF2 family, a group of MADS-box genes. Mef2D is expressed in many tissues, including neurons, muscle, adipose, and immune cells [7–10]. Mef2D regulates the expression of genes needed for cell survival and mitochondrial function in muscle and neurons [7–10]. Mef2D’s effect on mitochondrial function has been primarily shown to be through its transcriptional control of the mitochondrial DNA encoded mt-NADH dehydrogenase 6 (mtND6) [11, 12]. Given the effect of Mef2D on cell survival and mitochondrial function, an important question is how it affects the pancreatic beta-cell.

Human pancreatic imaging from the Human Protein Atlas shows clear pancreatic endocrine cell staining [13]. Only one study has investigated Mef2D activity in the beta-cell, demonstrating that Mef2D overexpression and knockdown modulate glucose-stimulated insulin secretion [14].

Given Mef2D expression in the endocrine pancreas, recently published data, and its apparent role in controlling mitochondrial function and cell survival, we sought to understand the function of Mef2D in the beta-cell. We confirmed Mef2D expression in INS-1 832/13 beta-cells and primary rat islets. Using lentiviral constructs, we built stable Mef2D overexpression and knockdown INS-1 832/13 beta-cell lines. Here, we present data on the effect of Mef2D overexpression and knockdown on mitochondrial function, cell survival, and insulin secretion.

## MATERIALS AND METHODS

### Cell culture

INS-1 832/13 beta-cells were maintained in complete RPMI 1640 medium with L-glutamine and 11.1 mM glucose supplemented with 50 U/ml penicillin, 50 μg/ml streptomycin, 10 mM HEPES, 10% fetal bovine serum, and INS-1 supplement (glutamine, Na-pyruvate, and 2-mercaptoethanol) [15].

### Lentivirus production and MEF2D manipulation

Rat shRNA constructs against Mef2D (5’-TTTCCGTGGCAACACCAAGTTTACTCAGC-3’) and a control 29-mer scrambled shRNA cassette (termed shCTRL) were cloned into the pGFP-C-shLenti plasmid, as were Lentivirus rat Mef2D overexpression particles and control GFP overexpression particles were produced and acquired from OriGene. INS-1 832/13 cells were transduced with the respective recombinant lentivirus at a multiplicity of infection of three viral particles/cell in the presence of polybrene (10mg/ml) for 48 hr. Following transduction, lentivirus transduced cells were selected through culture with puromycin (2 μg/ml), after which cells were transferred to normal culture media for experimental conditions.

### Glucose-stimulated insulin secretion

Glucose-stimulated insulin secretion (GSIS) was performed as described previously [16]. Briefly, cells were grown to confluency, washed with PBS, and preincubated in secretion assay buffer (SAB) containing 2.5 mM glucose for 2 h (114 mM NaCl, 4.7 mM KCl, 1.2 mM KH2PO4, 1.16 mM MgSO4, 20 mM HEPES, 2.5 mM CaCl2, and 0.2% BSA, pH 7.2). GSIS was performed by incubating quadruplicate replicate wells of cells in 1xSAB containing 2.5 mM glucose for 1 h, followed by 1 h in 1xSAB with 12 mM glucose, followed by collection of the respective buffers, as described previously. For total insulin content, cells were lysed in RIPA buffer with protease inhibitors (Life Technologies). Secreted insulin and total insulin were measured in 1xSAB using a rat insulin ELISA [16, 17].

### Oroboros respiration (O2k)

Experiments were done with permeabilized INS-1 832/13 beta-cells using a Clarke Oxygen electrode high-resolution respirometer (Oxygraph-2K; Oroboros Instruments, Innsbruck, Austria). For permeabilization, 832/13-derived cells were harvested in Miro5 respiration media (0.5 mM EGTA, 3 mM MgCl2, 60 mM K-lactobionate, 20 mM taurine, 10 mM KH2PO4, 20 mM HEPES, 110 mM sucrose, and 1 g/l BSA, pH 7.1) and permeabilized with digitonin (8 μM) [16]. All experiments were run in pairs with controls (NT, Lenti-shCTRL, Lenti-shMef2D, Lenti-GFP, orLenti-Mef2D). Respiration experiments were done with an oxygen concentration ranging between 200 and 450 nmol/ml at 37°C [16].

A basal, Complex I respiratory capacity was determined in the presences of malate (2 mM) and glutamate (10 mM). The respiratory capacity of Complex V and Complex II was assessed upon the addition of ADP (2.5 mM) and succinate (10 mM). Mitochondrial integrity was tested by adding cytochrome c (10 μM). Uncoupled maximal respiratory capacity of the electron transfer system was measured after a stepwise addition of 0.5 μM FCCP. Complex II mediated oxidative phosphorylation was assessed using Rotenone (0.5 μM). Residual oxygen consumption was determined following inhibition of Complex III with the addition of antimycin A. This state of residual oxygen consumption served as a baseline correction for all other states.

### Cell Viability Assays

INS-1 832/13 beta-cells untreated or transduced with Lenti-shMef2D, Lenti-Mef2D, Lenti-GFP, or Lenti-shCTRL were plated at a concentration of 2 × 10^5^ cells/ml in 96-well plates (at 100 μl/well). Cells were treated with etoposide (9 uM), camptothecin (2uM), or thapsigargin (0.3uM) and cell viability was determined using Alamar Blue assays (Sigma-Aldrich). Absorbance for Alamar Blue assays was read at 570 nm and 600 nm on a SpectraMax iD3 micro plate reader.

### Quantitative real-time polymerase chain reaction (qPCR)

RNA was harvested using TRI Reagent (Life Technologies), and cDNA was synthesized was completed using the High-Capacity cDNA Reverse Transcription kit (Life Technologies) [16]. Real-time PCRs were performed using the Life Technologies Biosystem 6 sequence detection system and software (Life Technologies). SYBR green-based primers were used to detect rat Nr4a1 (Forward 5’-GTGTTGATGTTCCTGCCTTTG-3’; Reverse 5’-TCAGACAGCTAGCAATGCG-3’), Mef2D (Forward 5’-TCATCTTCAACCACTCCAACA-3’; Reverse 5’-CGTTGAAACCCTTCTTCCTCA-3’), mtND6 (Forward 5’-ATCCCCGCAAACAATGACCA-3’; Reverse 5’-TTGGGGTTGCGGCTATTTAT-3’), Ndufv3 (Forward 5’-CAGGACCAACTAGCAAGACG-3’; Reverse 5’- CTAGGAAGGTGAATGCGTTGT-3’), oxogluterate dehydrogenase (Ogdh), Dihydrolipoamide S-succinyltransferase (Dlst), dihydrolipoamide dehydrogenase (Dld) (Forward 5’-GCAGGAGTAATTGGTGTGGAA-3’; Reverse 5’-CCTTGCCTCTGAAGTATACGTT-3’), Isocitrate dehydrogenase (Isdh) (Forward 5’-AACCGTGTGGCTCTAAAGG-3’; Reverse 5’-TCTTACAGTGGATGACGTTGG-3’), Malate dehydrogenase (Mdh2), Succinate dehydrogenase b (Sdh) (5’-ACAGTATCTGCAATCCATCGAG-3’; Reverse 5’- TGTCTCCGTTCCACCAGTA-3’), INS1+2 (5’-TGTCAAACAGCACCTTTGTGG-3’; Reverse 5’-GGGCCTCCACCCAGCTCCAGTT-3’), GLUT2, and Peptidylprolyl Isomerase A (PPIA) (Forward 5’-CCATTATGGCGTGTGAAGTC-3’; Reverse 5’- GCAGACAAAGTTCCAAAGACAG-3’) (internal control).

### Immunoblot analysis

Cells were lysed in radioimmunoprecipitation buffer supplemented with protease and phosphatase inhibitors (Pierce). Total cellular protein was quantified with the BCA protein assay kit (Pierce) [15, 16]. Protein samples (30 μg) were heated to 70°C in Laemmeli buffer with β-mercaptoethanol and resolved on a 10% PAGE gel. Proteins were transferred to PVDF, and membranes were blocked for 1 hour in PBS-Blocking buffer, followed by overnight incubation in the appropriate antibody. The following antibodies were used: tubulin (1:6,000; Proteintech), Mef2D (1:1,000; BD Biosciences), mtND6 (1:500; Genetex), OxPHOS (1:1,000; Invitrogen), GLUT2 (1:500; Genetex). Primary antibodies were detected by incubating with the appropriate secondary antibody for 1 hour (Licor). Bands were detected by imaging on Odyssey Clx with image studio software and quantified using ImageJ.

### Statistical analysis

All treatment experiments were performed in at least triplet, and analysis will be reported as means ± standard deviation. For analysis between 2 groups, the student’s t-test was used. One-way analysis of variance (ANOVA) was used for comparing 3 or more groups. Two-way ANOVA was used for comparing 3 or more groups with 2 or more treatments. All statistics were performed in GraphPad Prism with p<0.05 considered significant.

## RESULTS

### Mef2D overexpression or knockdown affects mitochondrial respiration in the beta-cell

Our goal is to understand the effect of Mef2D overexpression and knockdown in the beta-cell. To do this we developed lentiviral constructs to produce INS-1 832/13 Mef2D overexpression and knockdown cell lines. We validated that Mef2D overexpression significantly increased Mef2D mRNA and protein levels (Figure 1A-E). An additional band is seen in our Mef2D overexpression, indicating a splice variant of Mef2D is produced by the lentiviral Mef2D overexpression line (Figure 1D-E). Similarly, Mef2D knockdown significantly decreased Mef2D mRNA and protein levels (Figure 1F-H). These tools allow us to test the function of Mef2D in the beta-cell.

**Figure 1.**
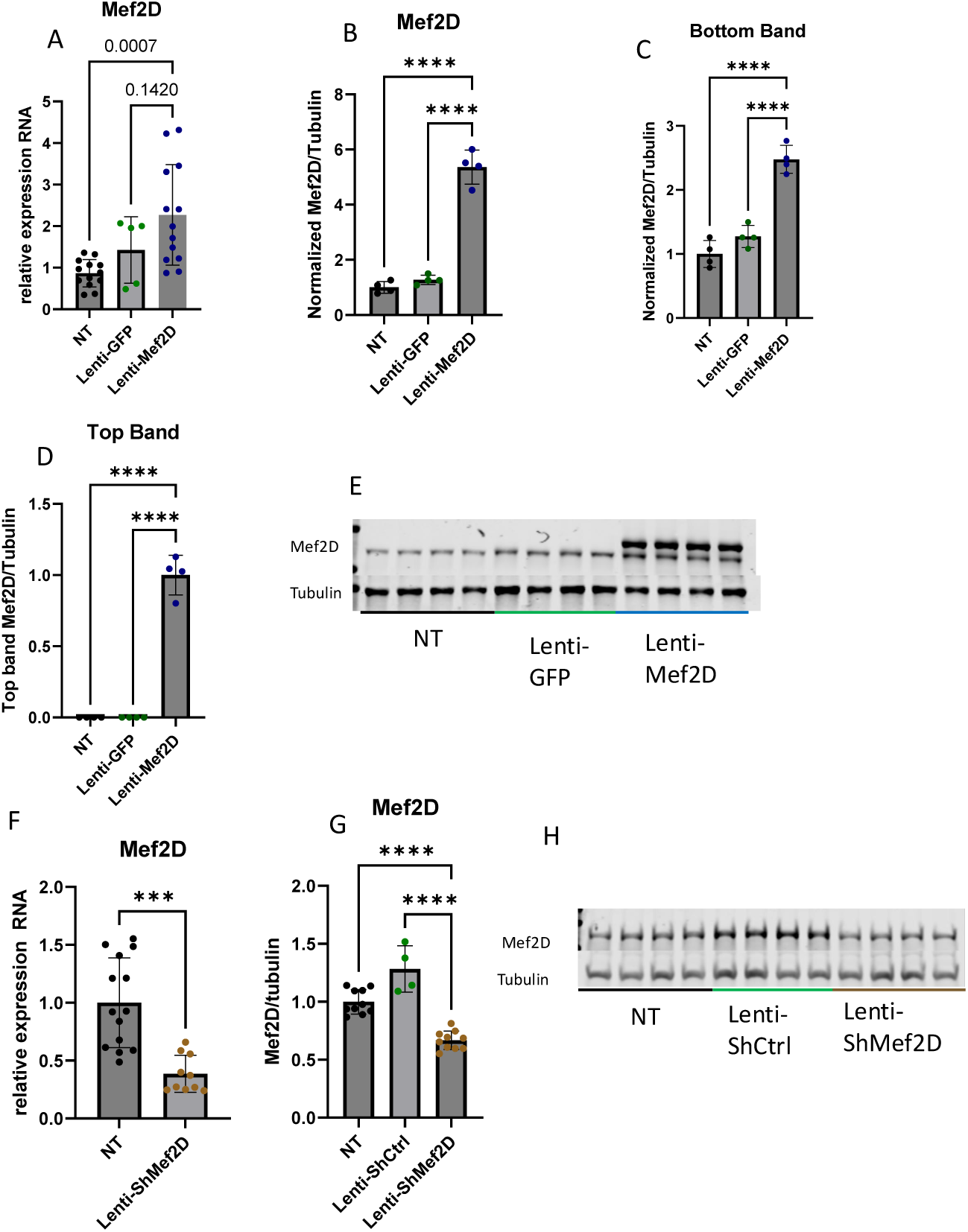
Mef2D overexpression or knockdown in the INS-1 832/13 beta-cell. A-E) Confirmation of Lenti-Mef2D overexpression line using mRNA and protein. Top band and bottom band of samples quantified. F-H) Confirmation of Lenti-ShMef2D knockdown line using mRNA and protein. p-values *<0.05, **<0.01, ***<0.001, or ****<0.0001.

In neurons, Mef2D induces the expression of mitochondrial genes that are components of the electron transport chain (ETC) Complex I. Given these findings, we hypothesized that Mef2D overexpression and knockdown would impinge on mitochondrial respiration. Cells overexpressing Mef2D had impaired Complex I, Complex II, Complex V, and Complex II linked oxidative phosphorylation and maximal respiration (Figure 2A-E). Cells deficient in Mef2D had increased respiration only in Complex II linked oxidative phosphorylation and maximal respiration (Figure 2F-J). The parental INS-1 832/13 beta-cells had no difference in respiration compared to the Lenti-shCTRL cell lines, demonstrating that lentiviral transduction does not impair respiration (Figure 2K-O). This data indicates that Mef2D overexpression strongly impairs beta-cell mitochondrial respiration, while Mef2D knockdown modestly enhances respiration at the oxidative phosphorylation and uncoupled respiration steps in the beta-cell.

**Figure 2.**
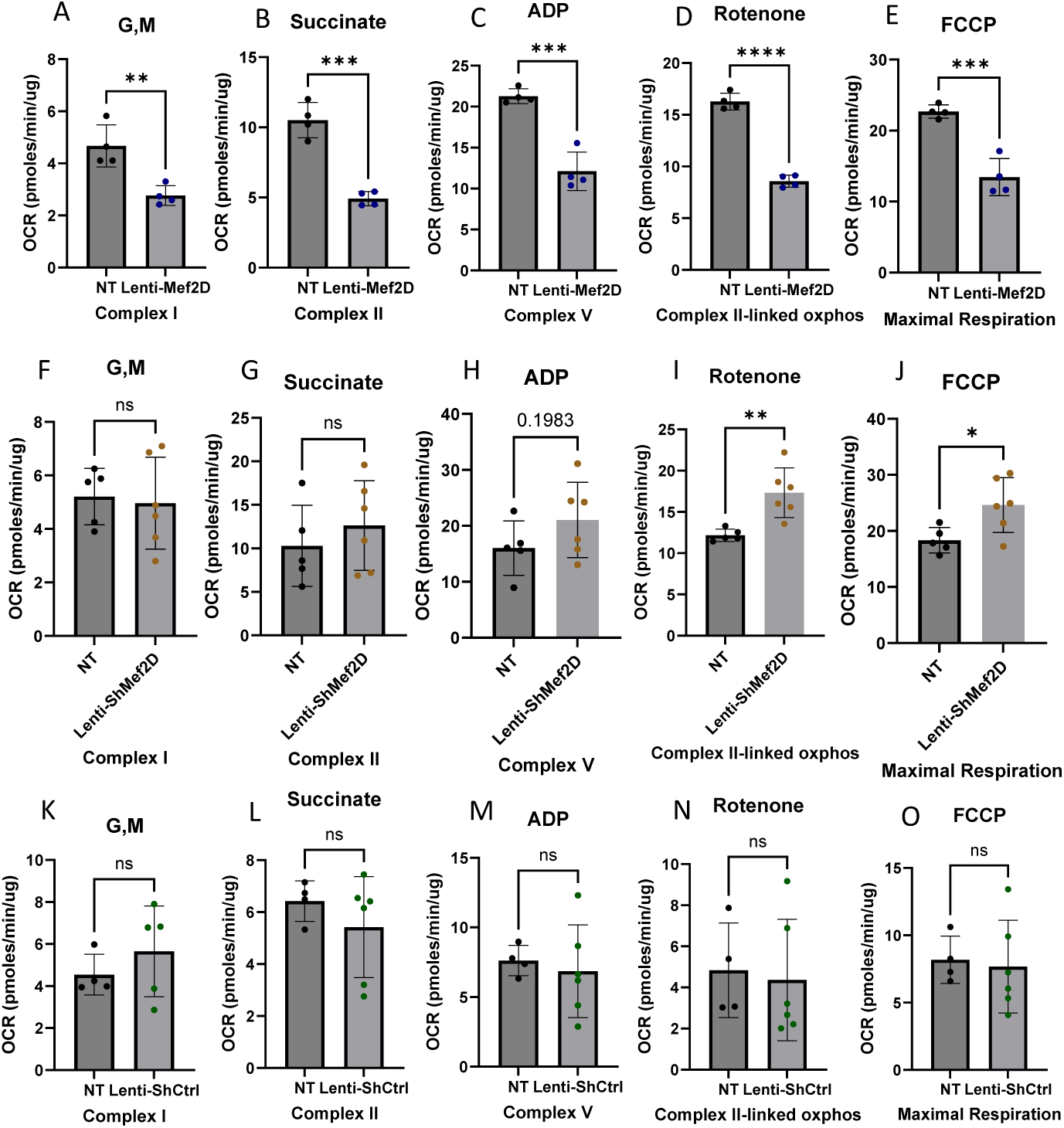
Mef2D overexpression or knockdown affects mitochondrial respiration in the beta-cell. Oxidative phosphorylation was measured in both Mef2D overexpression and knockdown lines. A, F) Glutamate and Malate (G, M) used to stimulate Complex I. B, G) Succinate to stimulate Complex II. C, H) ADP is used to stimulate Complex V. D, I) Rotenone is used to inhibit Complex I activity to allow data to be gathered for Complex II linked oxidative phosphorylation. E, J) FCCP is a proton uncoupler to allow for measurement of the maximal respiration of the cells. K-O) No treatment was compared with Lenti-ShCtrl across the steps to confirm no change with lentiviral induction. p-values *<0.05, **<0.01, ***<0.001, or ****<0.0001.

### Mef2D overexpression results in decreased expression of electron transport chain components in the beta-cell

Given the observed changes in respiration with Mef2D overexpression, we sought to determine the mechanism for impaired respiration. Based on neuronal data demonstrating that Mef2D regulates components of the electron transport chain, we measured the expression of various components in cells with Mef2D overexpression. We measured the expression of the Complex I components Ndufv3 and mtND6. We saw decreased mRNA expression of Ndufv3 with Mef2D overexpression, with no changes in Ndufv3 levels with Mef2D knockdown (Figure 3A-B). We also observed decreased mRNA and protein levels of mtND6 with Mef2D overexpression (Figure 3C-E). Finally, we measured protein levels of various electron transport chain components and saw decreases for Complex I (Ndufb8) protein expression and a significant decrease in Complex II (SDHB) protein levels (Figure 3F-J). These data demonstrate that Mef2D overexpression in the beta-cell decreases the expression of electron transport chain components.

**Figure 3.**
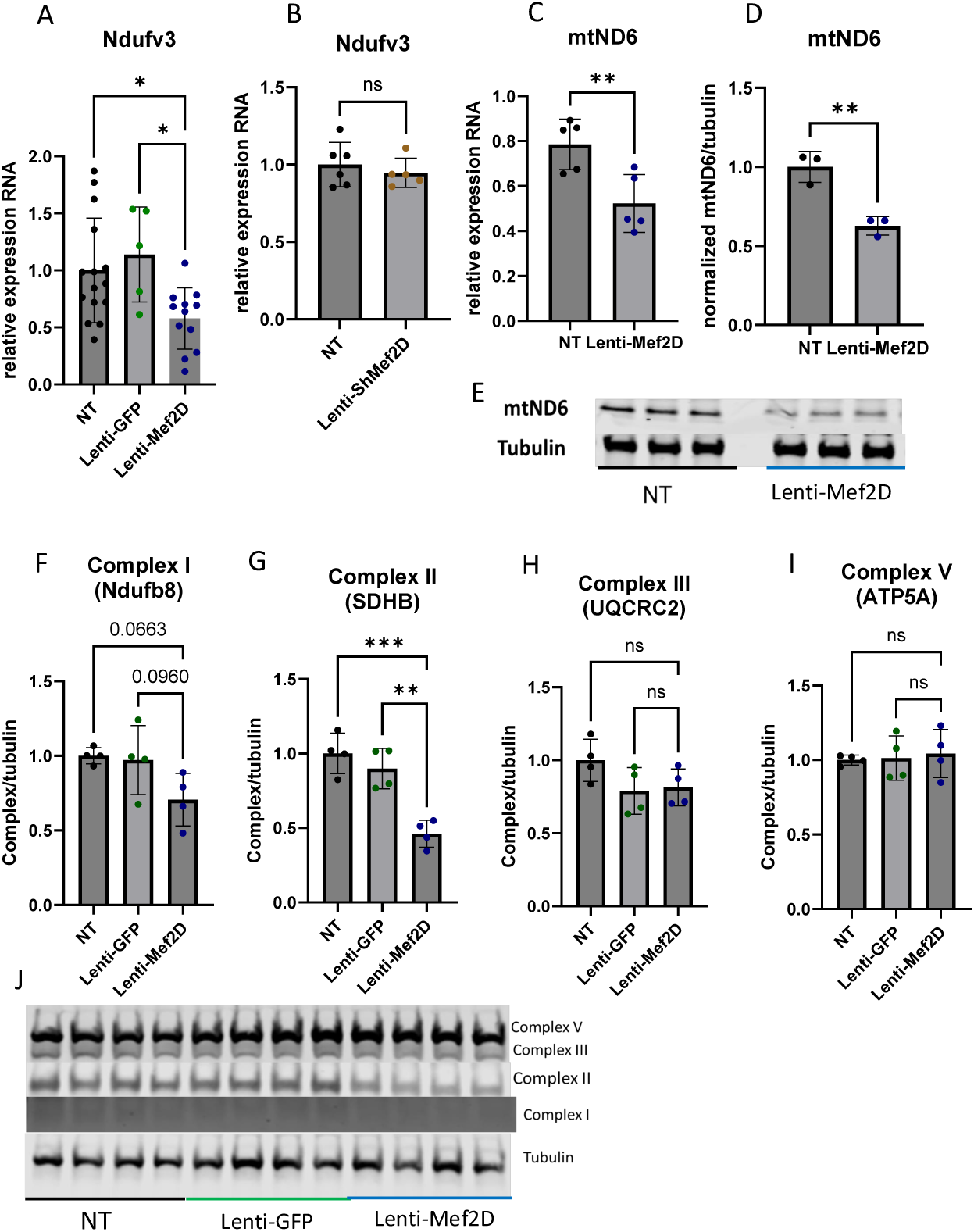
Mef2D overexpression results in decreased expression of electron transport chain components in the beta-cell. A, B) mRNA measurement of Ndufv3 in Lenti-Mef2D and Lenti-ShMef2D. C-E) Protein and mRNA measurement of mtND6 in Lenti-Mef2D. F-J) Protein measurement of electron transport chain components Complex I, II, III, and V. p-values *<0.05, **<0.01, ***<0.001, or ****<0.0001.

### Mef2D overexpression in the beta-cell results in decreased levels of the TCA dehydrogenase component Ogdh

The electron transport chain functions due to the shunting of NADH and FADH2 into complex I and II, respectively, from TCA cycle dehydrogenases. We hypothesize that decreased TCA cycle dehydrogenases caused inhibition of mitochondrial function. To test this hypothesis, we measured the expression of TCA cycle dehydrogenases. In cells with Mef2D overexpression, we measured the expression of Mdh2, Isdh, Sdh, Dld, and Dlst and saw no significant change in expression (Figure 4A-E). In addition, we measured the expression of Ogdh, a component of the alpha-ketoglutarate dehydrogenase complex, and observed a significant decrease in its expression (Figure 4F). However, cells deficient in Mef2D showed no changes in any of the TCA cycle dehydrogenases (Figure 4G-L). These data demonstrate that overexpression of Mef2D impairs the TCA cycle through downregulating Ogdh, a critical component of the TCA cycle.

**Figure 4.**
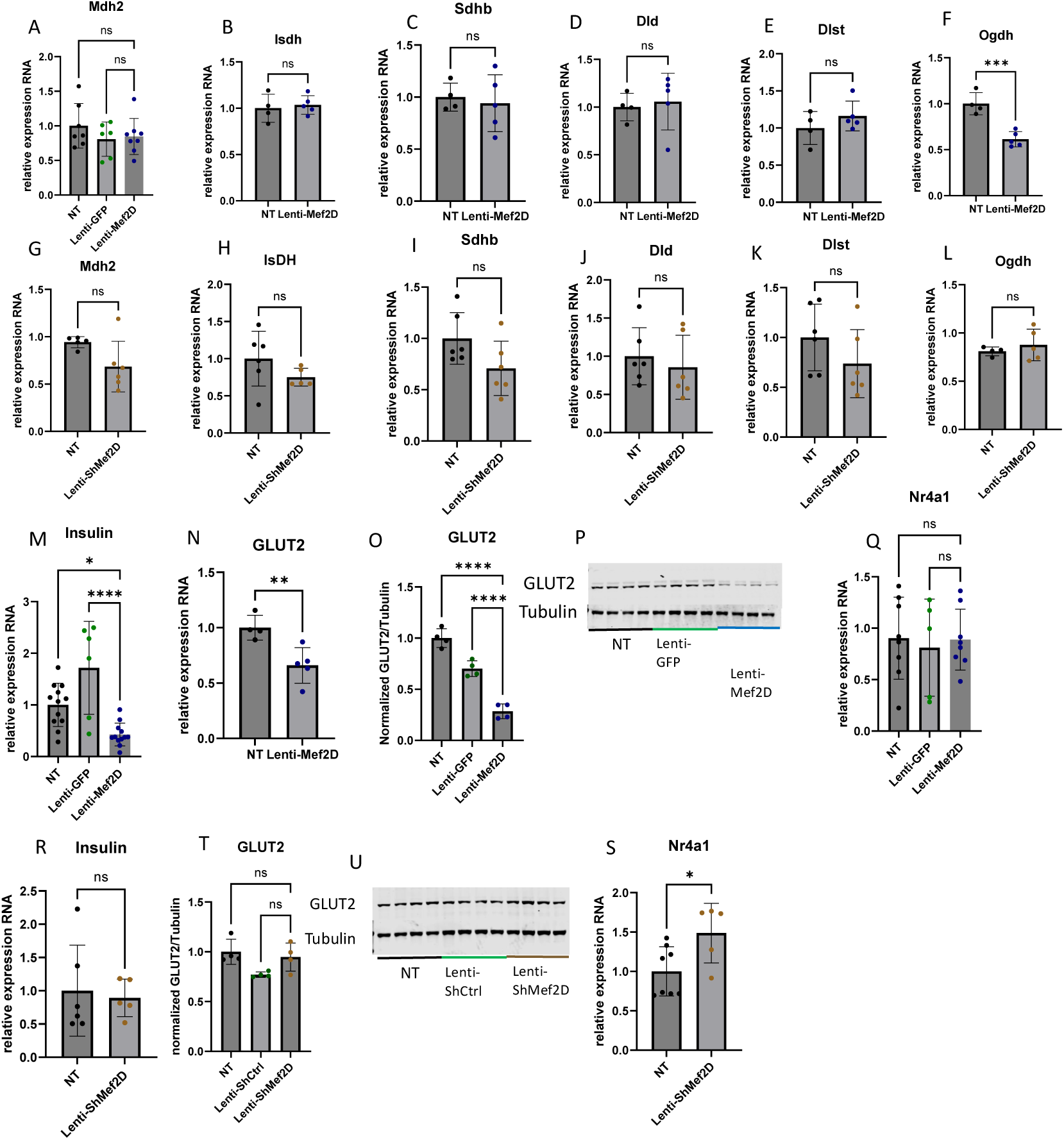
Mef2D overexpression in the beta-cell results in decreased levels of the TCA dehydrogenase component Ogdh. A-F) mRNA measurement of dehydrogenases in TCA and ETC of Lenti-Mef2D. G-L) mRNA measurement of dehydrogenases in TCA and ETC of Lenti-ShMef2D. M,R) mRNA measurement of Insulin in Lenti-Mef2D and Lenti-ShMef2D respectively. N-P) mRNA and protein measurement of GLUT2 in Lenti-Mef2D. T, U) Protein measurement of GLUT2 in Lenti-ShMef2D. Q, S) mRNA measurement of Nr4a1 in Lenti-Mef2D and Lenti-ShMef2D respectively. p-values *<0.05, **<0.01, ***<0.001, or ****<0.0001.

We hypothesize that in addition to altering expression of genes involved in the TCA cycle and the ETC, that changes in Mef2D levels will alter expression of critical beta-cell genes. Cells overexpressing Mef2D showed decreased insulin mRNA levels and decreased GLUT2 mRNA and GLUT2 protein levels (Figure 4M-P). Conversely, cells deficient for Mef2D showed no change in insulin mRNA and no change in GLUT2 protein expression (Figure 4R-U). Cells deficient in Mef2D did show an increase in Nr4a1 mRNA expression, while overexpression of Mef2D did not change Nr4a1 mRNA expression (Figure 4Q, S). These data demonstrate that Mef2D overexpression impairs key genes involved in beta-cell function.

### Mef2D modulates Glucose-stimulated Insulin Secretion

Beta-cell insulin secretion is directly tied to glucose metabolism and ATP production. Anything that modulates ATP production has the capacity to affect insulin secretion. Given the effects on mitochondrial respiration and expression of critical portions of the TCA cycle and ETC, we hypothesized that Mef2D overexpression would impair GSIS while knockdown would enhance GSIS. As anticipated, cells overexpressing Mef2D showed a significant reduction in GSIS (Figure 5A) with no change in total insulin content (Figure 5C). Conversely, cells deficient for Mef2D had increased unstimulated insulin release and GSIS (Figure 5B) with a significant increase in total insulin content (Figure 5D). These data demonstrate that Mef2D overexpression impairs GSIS, while Mef2D knockdown enhances GSIS.

**Figure 5.**
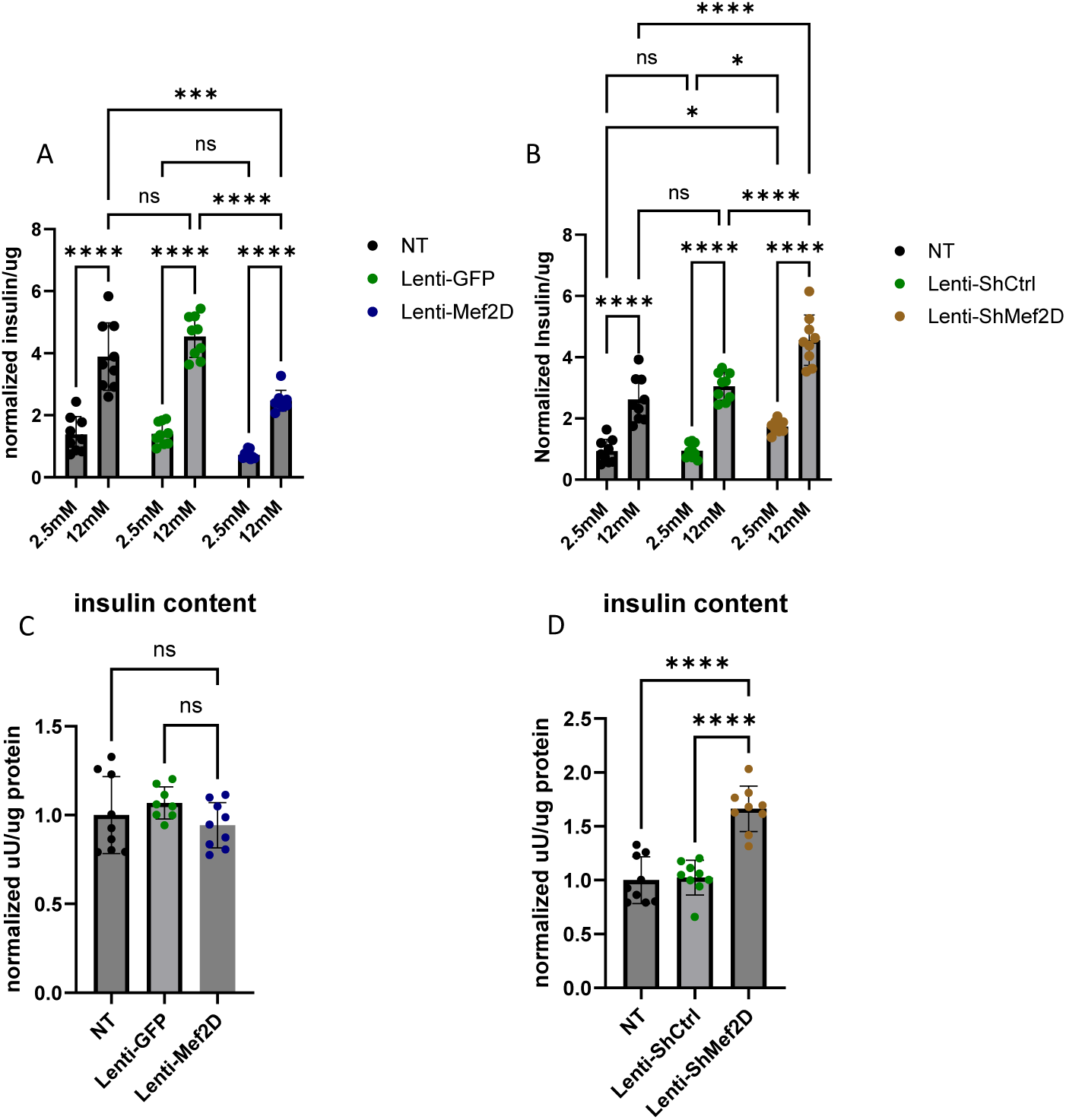
Mef2D modulates glucose-stimulated insulin secretion. A) Glucose stimulated insulin secretion in Lenti-Mef2D line is inhibited. B) Insulin content in Lenti-Mef2D. C) Glucose stimulated insulin secretion in Lenti-ShMef2D is enhanced. D) Insulin content in Lenti-ShMef2D is enhanced. p-values *<0.05, **<0.01, ***<0.001, or ****<0.0001.

### Mef2D impairs Cell Viability

Mitochondrial dysfunction can directly impair cell viability. We showed changes to mitochondrial respiration and mitochondrial gene expression due to Mef2D overexpression or knockdown. We hypothesized that Mef2D-mediated mitochondrial defects may cause beta-cells to be more susceptible to cell damage and impair cell survival. We measured viability by Alamar Blue assay in cells overexpressing or deficient for Mef2D. Mef2D overexpression decreased overall cell viability. Additionally, viability was decreased in Mef2D overexpressing cells with etoposide, camptothecin, or thapsigargin treatment (Figure 6A). Conversely, cells deficient for Mef2D had no change in viability under standard culture conditions or with camptothecin treatment; however, there was increased cell viability under etoposide or thapsigargin treatment (Figure 6B). These data demonstrate that under stressed conditions, Mef2D overexpression decreases cell viability and that Mef2D reduction will rescue cell viability.

**Figure 6.**
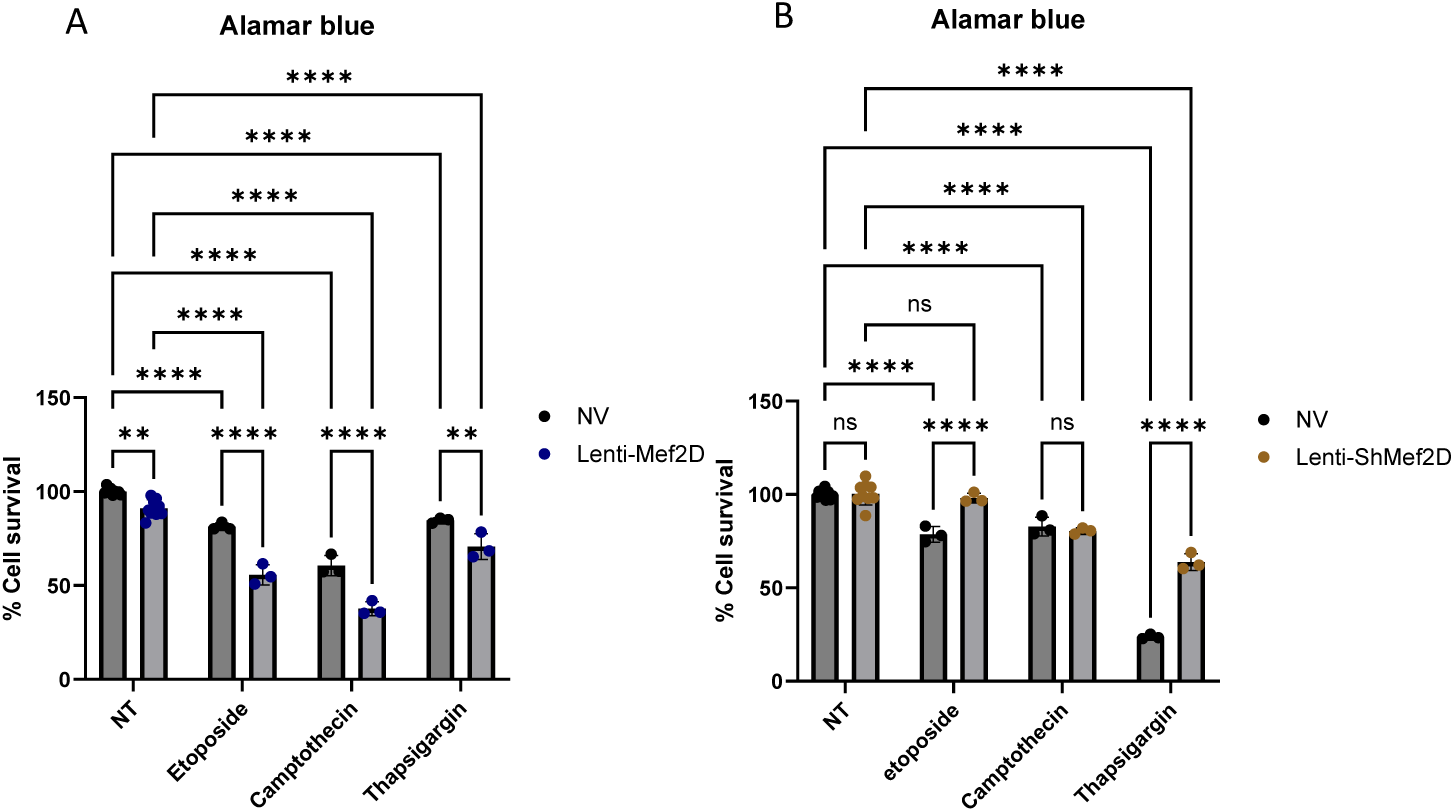
Mef2D overexpression impairs cell viability. A) Alamar blue measurement of cell viability in Lenti-Mef2D treated with Etoposide, Camptothecin, Thapsigargin. B) Alamar blue measurement of cell viability in Lenti-ShMef2D treated with Etoposide, Camptothecin, Thapsigargin. p-values *<0.05, **<0.01, ***<0.001, or ****<0.0001.

## DISCUSSION

The pancreatic beta-cell plays a crucial role in type 1 and type 2 diabetes. Both types display a loss of beta-cells and decreased glucose tolerance. Given the high expression of Mef2D in the human beta-cell, as presented by the Human Protein Atlas [13], and recent publications showing that Mef2D has an inhibitory effect on glucose-stimulated insulin secretion in the beta-cell [14], we sought to further explore its function. Similar to the previous published findings, we validated that knock down of Mef2D resulted in increased insulin secretion (Figure 5C) [14]. In support of this inhibitory effect on insulin secretion, cells with Mef2D overexpression showed inhibited glucose-stimulated insulin secretion (Figure 5A). As the previous study did not explore the mechanism by which Mef2D expression modulated glucose-stimulated secretion, we investigated the different pathways and genes involved in beta-cell insulin secretion under Mef2D knockdown and overexpression.

The mitochondria are essential for insulin secretion. Given the changes we observed in insulin secretion, we measured mitochondrial expression in the presences of Mef2D overexpression and Mef2D knockdown. In cells with Mef2D overexpression, mitochondrial respiration is inhibited across all measured complexes and in maximal respiration (Figure 2A-E). A reduction in Complex I components Ndufv3 and mtND6 and a reduction in Complex II (SDHB) expression would lead to reduced respiration (Figure 3A, C-E, G, J). In neurons, a decline of mtND6 led to reduced Complex I expression and activity [11]. However, in neurons, a knockdown of Mef2D resulted in the reduction of mtND6, and conversely, Mef2D overexpression in beta-cells leads to a reduction of mtND6 (Figure 3C-E) [11]. While Mef2D overexpression does not affect Complex V expression, the reduced function is likely due to the decreased expression and function of Complex I and Complex II, which would result in a reduced proton gradient. The decreased proton gradient also likely results in higher levels of oxidative stress from the reduced usage of the H+, leading to the creation of reactive oxygen species, which also lowers insulin secretion [18]. As the TCA cycle will feed NADH and FADH2 into the ETC, we also looked at some of the TCA cycle dehydrogenases. Cells with Mef2D overexpression had reduced Ogdh expression, a component of the alpha-ketoglutarate dehydrogenase complex (Figure 4F). A reduced function of Ogdh would result in lower NADH production, NADH is oxidized by Complex I, and reducing acetyl-Coa catabolism, possibly impinging on earlier steps in the TCA cycle. The Mef2D knockdown line showed increased maximal respiration (Figure 2J) and increased Complex II-linked oxidative phosphorylation, indicating the direct link to the TCA cycle as Complex II is also succinate dehydrogenase (Figure 2I). The dehydrogenases and Ndufv3 were measured, and there were no changes in expression (Figure 4B & 5G-L). This data indicates that one way by which Mef2D inhibits glucose-stimulated insulin secretion is through the inhibition of mitochondrial respiration by targeting components of the ETC and reducing Ogdh in the TCA cycle. However, the mechanism by which Mef2D knockdown increased insulin is only partially explained with some increased respiration. Given this, further measurements of genes within the ETC may yield more answers.

Another critical piece of insulin secretion is glucose sensing. In the beta-cell, glucose primarily enters through GLUT2 [19]. In the Mef2D overexpression line, GLUT2 expression is reduced, which will lead to reduced glucose uptake. Decreased glucose uptake will impinge on the glucose-stimulated insulin secretion at the initiatory stages (Figure 4N-P). Mef2D overexpression reduced insulin mRNA, but insulin content was unchanged (Figure 4M & 5C). Conversely, cells with Mef2D knockdown showed no change of GLUT2 expression (Figure 4T, U). Mef2D knockdown did not change insulin mRNA, however insulin content was increased (Figure 4R & 5D). What is clear is that changes in Mef2D levels significantly affect multiple aspects of the insulin secretion pathway, from glucose entry to insulin production and secretion. To completely understand the role of Mef2D in these phenotypic aspects, further experiments will be needed.

There is a reduction of beta-cell cell viability in both major forms of diabetes. In neurons, Mef2D has been shown to play a critical role in maintain cell viability [20–22]. The Mef2D overexpression line shows reduced cell viability in standard culture and under pharmacologically stressed conditions (Figure 6A). This reduced cell viability is likely due to reduced mitochondrial respiration (Figure 2A-E). The lowered proton gradient from reduced Complex I and II activity would likely result in higher levels of oxidative stress, which would reduce cell viability. There is also a link between Complex I components, such as mtND6, and reduced cell viability [11, 23]. Our data demonstrates that mtND6 and the Complex I component Ndufv3 were reduced in cells with Mef2D overexpression, which supports the published literature (Figure 3A, C-E).

While the Mef2D overexpression line showed decreased cell viability, the Mef2D knockdown line shows increased cell viability under pharmacologically stressed conditions (Figure 6B). The increase in cell viability could be due to increased mitochondrial respiration (Figure 2I, J). The gene Nr4a1 has been linked to beta-cell mitochondrial respiration, and Mef2D has been shown to control Nr4a1 transcription in neurons and T-cells [16, 20, 24, 25]. Nr4a1 induction leads to improved cell viability in neurons, and induces apoptosis in T-cells, demonstrating tissue specific effects [20, 26]. The Mef2D overexpression line does not affect Nr4a1 expression (Figure 4Q).

However, the Mef2D knockdown line demonstrates increased Nr4a1 expression (Figure 4S). Induced Nr4a1 expression in Mef2D knockdown lines may be essential for the increased observed cell viability. Future studies will need to be concluded to determine if Nr4a1 is a direct Mef2D target, and if Mef2D knockdown in beta-cells lacking Nr4a1 lose the observed increased survival phenotype.

## CONCLUSION

Our study indicates that Mef2D has a regulatory role in the beta-cell insulin secretion and cell viability pathways (Figure 7). Mef2D overexpression inhibits insulin secretion through reduced mitochondrial respiration. Decreased ETC and TCA cycle component expression will lead to reduced mitochondrial respiration. Mef2D overexpression reduced GLUT2 expression, reducing glucose uptake and inhibiting glucose sensing in the beta-cell. Mef2D overexpression also reduces cell viability, likely through the reduced mitochondrial function. Conversely, Mef2D knockdown shows increased insulin secretion through some enhancement in mitochondrial respiration and increased insulin content. Mef2D knockdown also shows increased cell viability, which stems from the increased mitochondrial respiration and increased Nr4a1 expression. Our study shows a more in-depth and unique view of Mef2D’s roles within the beta-cell. These findings demonstrate a new target that maybe utilized to increase functional beta-cell mass.

**Figure 7.**
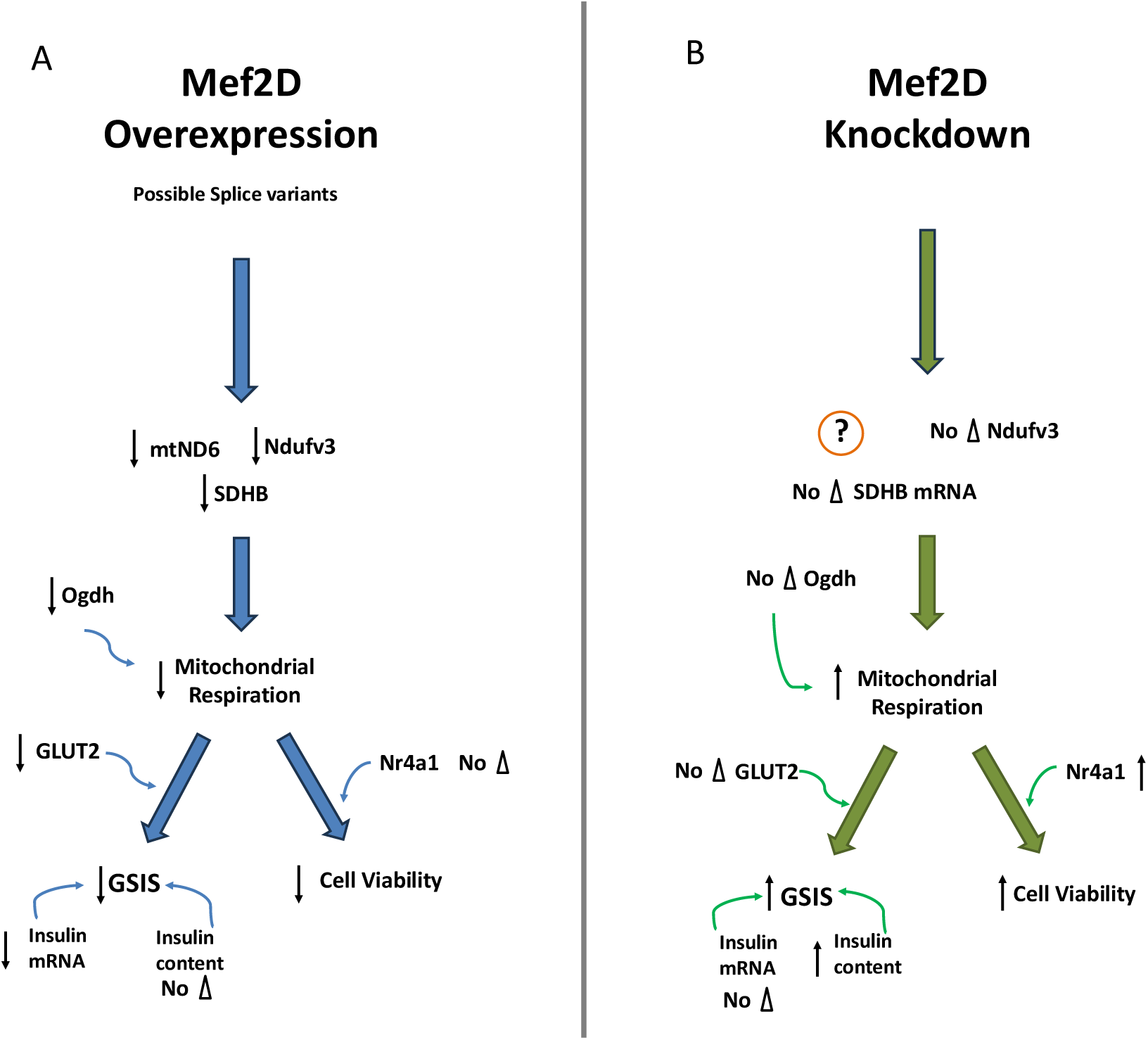
Mef2D will affect GSIS and cell viability within the beta-cell. A) Mef2D overexpression will reduce the expression of the ETC components mtND6, Ndufv3, and SDHB, reducing mitochondrial respiration. The TCA dehydrogenase Ogdh will add to the reduced mitochondrial respiration. Reduced mitochondrial respiration, along with reduced GLUT2 expression, will lead to reduced GSIS. Reduced insulin mRNA will add to the reduced GSIS. Mitochondrial respiration is also leading to reduced cell viability. B) Mef2D knockdown leads to enhanced mitochondrial respiration and increased insulin content, leading to increased GSIS. The increased mitochondrial respiration, along with increased Nr4a1, will also lead to increased cell viability.

## ACKNOWLEDGMENTS

We thank members of the Sharma research group for their productive discussions that have assisted with this work. This work was supported by a grant from the National Institutes of Health R15DK144832 to JST.

## AUTHOR CONTRIBUTIONS

Conceptualization: J.E.C, R.B.S, and J.S.T. ; Experimental Design and Performance: J.E.C and J.S.T; Data Analysis: J.E.C. and J.S.T.; Writing-original draft: J.E.C. and J.S.T.; Writing-review and editing: J.E.C, R.B.S, and J.S.T.; Supervision: J.S.T.; Funding Acquisition: J.S.T.

### DECLARATION OF INTEREST

The authors have no declarations of interests.

## Notes

### Competing Interest Statement

The authors have declared no competing interest.

